# Adenovirus-α-defensin complexes induce NLRP3-associated maturation of human phagocytes via TLR4 engagement

**DOI:** 10.1101/2021.10.26.465098

**Authors:** Karsten Eichholz, Tuan Hiep Tran, Coraline Chéneau, Thi Thu Phuong Tran, Océane Paris, Martine Pugniere, Eric J Kremer

**Author notes:** Equal contribution (Karsten Eichholz, Tuan Hiep Tran, Coraline Chéneau). Faculty of Pharmacy, PHENIKAA University, Yen Nghia, Ha Dong, Hanoi 12116, Vietnam, PHENIKAA Research and Technology Institute (PRATI), A&A Green Phoenix Group JSC, No.167 Hoang Ngan, Trung Hoa, Cau Giay, Hanoi 11313, Vietnam. University of Science and Technology of Hanoi Vietnam Academy of Science and Technology, Department of Life Sciences, 18 Hoang Quoc Viet, Cau Giay, Hanoi, Vietnam (Thi Thu Phuong Tran).

## Abstract

Intramuscular delivery of human adenovirus (HAdV)-based vaccines leads to rapid recruitment of neutrophils, which then release antimicrobial peptides/proteins (AMPs). How these AMPs influence vaccine efficacy over the subsequent 24 hr is poorly understood. In this study, we asked if human neutrophil protein 1 (HNP-1), an α-defensin that influences the direct and indirect innate immune responses to a range of pathogens, impacts the innate response of human phagocytes to three HAdV species/types (HAdV-C5, -D26, -B35). We show that HNP-1 binds to the capsids, redirects HAdV-C5, -D26, -B35 to Toll-like receptor 4 (TLR4), which leads to internalization, an NLRP3-mediated inflammasome response, and IL-1β release. Surprisingly, IL-1β release was not associated with notable disruption of plasma membrane integrity. These data further our understanding of HAdV innate immunogenicity and may provide pathways to extend HAdV-based vaccines efficacy.

## Introduction

The immunogenicity of adenoviruses (AdVs) in the context of infection of immune suppressed individuals, therapeutic gene transfer, and vaccines is a pressing issue. With the occurrence of SARS-CoV-2, AdV-based (human and simian) vaccines are being administered on a global scale (1). Human AdVs (HAdVs) are ~950 Å diameter, nonenveloped proteinaceous particles containing a ~36,000 (± 9,000) double-stranded linear DNA genome. There are currently 7 HAdV species (A-G) and >100 types based on serology and phylogeny. HAdVs typically cause self-limiting respiratory, ocular, or gastro-intestinal tract infections in all populations regardless of health standards, but infections can be lethal in immune-compromised individuals (2). Due to their endemic nature at military training facilities, live HAdV types 4 and 7 have been used as vaccines since the 1950s to prevent severe respiratory illness in recruits (3). AdV-based vaccines have also been approved in a prime/boost regimen against Ebola virus in the EU and have been tested in more than 200 clinical trials (clinicaltrials.gov, May 2021).

In the context of natural infections, gene transfer, or vaccination, neutrophils and professional antigen-presenting cells (APCs) such as monocytes and dendritic cells (DCs) extravasate to the site of injection or infection and detect danger-associated molecular patterns (DAMPs) or pathogen-associated molecular patterns (PAMPs). APCs differentiate between pathogens via their pattern recognition receptors on the cell surface, in endosomal compartments, or in the cytosol. PRRs include Toll-like receptors (TLRs) (among which the most thoroughly studied receptor is TLR4), the nucleotide-binding oligomerization domain (NOD)-leucine-rich repeats (LRR)-containing receptors (NLR) (among which NLRP3 is the most thoroughly studied receptor to induce the inflammasome), the retinoic acid-inducible gene 1-like receptors (RIG-1), and the C-type lectin receptors. PRRs detect different PAMPs and DAMPs and hence, differentially affect the immune environment that is made up of pro-inflammatory cytokines, chemokines and host defence proteins/peptides (HDPs). In some inflammatory environments, monocytes mature into DCs and then migrate to lymphoid tissues to initiate antigen-specific T- and B-cell responses.

Antimicrobial peptides (AMPs) are effector molecules of the innate immune system, and act via antibiotic-like properties against a broad array of infectious agents (4, 5), or by promoting the activation and maturation of some APCs. Neutrophils are a rich source of AMPs: ~20% of their cytoplasmic content can be AMPs (6). The rapid release of HDPs acts as part of first-line responders to the disruption of tissue homeostasis (7). Lactoferrin, α-defensin, and cathelicidin (LL-37) are alarmins that also modulate innate and adaptive immune responses by directly engaging pattern recognition receptors in antigen-presenting cells (4, 5). Additionally, in epithelial cells, α-defensins impair HAdV-C5, -A12 and -B35 infection by stabilizing an intrinsically disordered region of the capsid vertex and thus prevents intracellular viral disassembly (8–10).

We recently reported how lactoferrin retargets HAdV-C5, -D26 and -B35 to TLR4 and initiates inflammasome assembly and interleukin 1 beta (IL-1β) release without loss of membrane integrity (11). The inflammasome, a multiprotein platform consisting of a pattern recognition receptor that promotes aggregation of apoptosis-associated speck-like protein containing a CARD, recruits and auto-activates pro-caspase 1. Pro-caspase-1 auto-activation can be followed by removal of the N-terminus of gasdermin D (GSDMD), which initiates the loss of plasma membrane integrity via pore formation (12). Canonical and non-canonical NLRP3 inflammasome formation is preceded by transcriptional priming event needed to produce inflammasome components and cytokines (13). In primary human monocytes, LPS-TLR4 engagement can induce an alternative inflammasome activation that is characterized by IL-1 β secretion in the absence of a transcriptional priming event, inflammasome aggregation, and pyroptotic cell death (14).

Here, we expanded our understanding of how an α-defensin impacts innate immune sensing of HAdV-C5, - D26 and -B35, which were chosen based on their development as vaccines (15–22). Mechanistically, HNP-1 binds HAdV-C5, -D26, and -B35 with affinities in the nanomolar range, increases HAdV uptake via a retargeting to TLR4 complexes, which in turn induces NLRP3 inflammasome formation and release of IL-1β.

## Results

### HNP-1 binds to HAdV-C5, -D26 and -B35 and increases infection of monocyte-derived DCs

HNP-1 binds the HAdV-C5 capsid through electrostatic and hydrophobic interactions at the juxtaposition of the penton base and fibre, and on hexon (8, 23). These interactions can reduce HAdV-C5 and HAdV-B35 infection of A549 cells (transformed human lung epithelial cells) [7]. To better understand the interactions, we quantified the affinity of human HNP-1 to HAdV-C5, -D26 and -B35by surface plasmon resonance (SPR) analyses. We show that HNP-1 binds to HAdV-C5, -D26 and -B35 with KDs that varied from 37 to 76 nM (**Figure 1A & B**) with a two-state association-dissociation reaction (**Figure S1A**).

**Figure 1).**
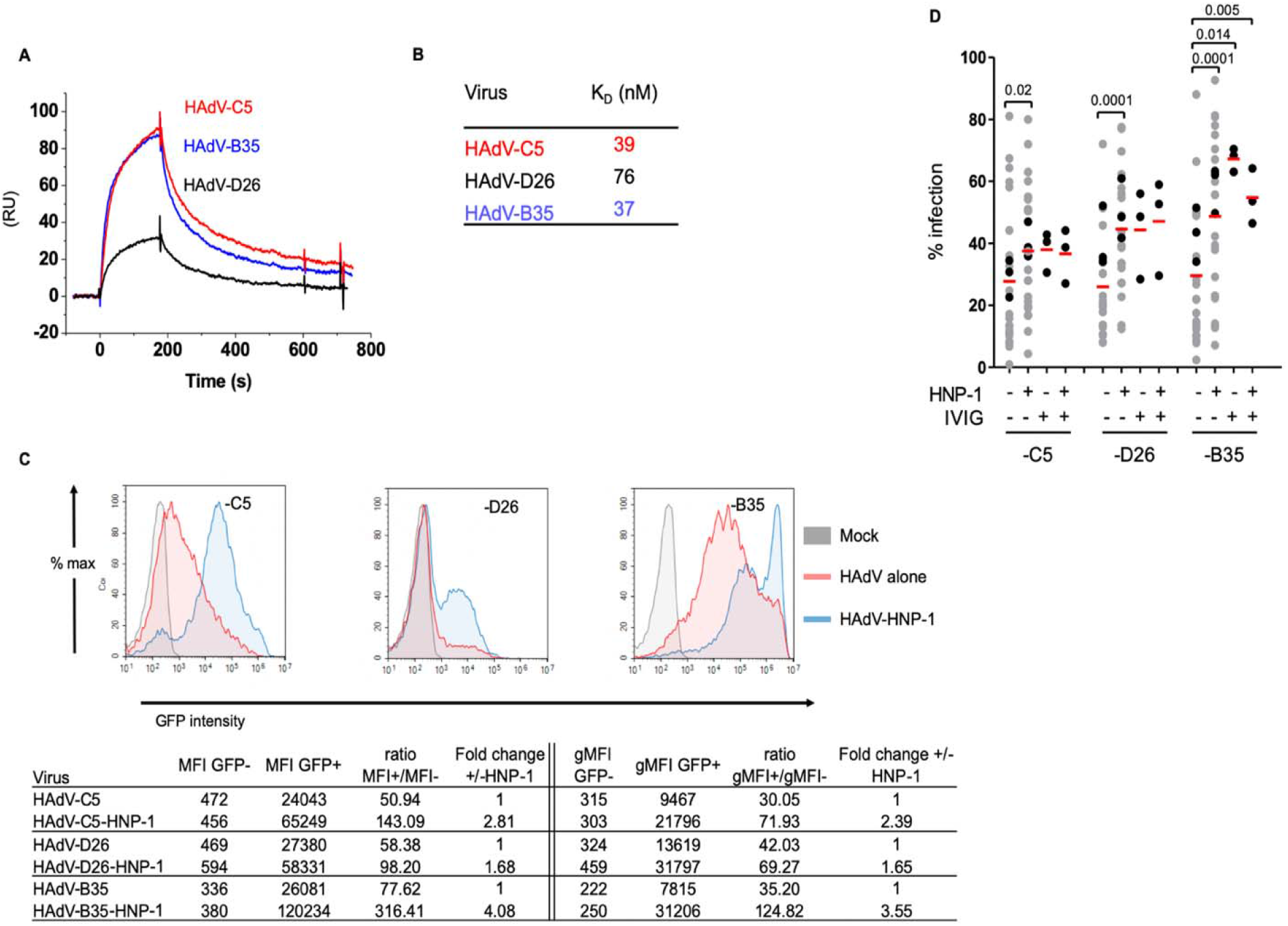
HNP-1 interacts with HAdVs and increase infection of DCs. **A)** SPR sensorgrams of HNP-1 - HAdV interactions: HAdV-C5 (red), -D26 (black), and -B35 (blue) were coupled to the sensor chip. A range of 6.25 - 200 nM of HNP-1 was injected and the KD was calculated (RU = resonance units). **B)** KD of HNP-1 for HAdV-C5, -D26 and -B35. **C)** Representative profiles of DCs infected with HAdV-**C5**, -D26, and -B35. Mock-treated DCs (grey), or incubated with HAdV-C5, -D26 or -B35 + HNP-1. MFI and gMFI for the FP+ and FP-cells for the different vectors ± HNP-1 and the fold changes in presence of HNP-1 for a representative experiment are shown. **D)** Cumulative data using HAdVs complexed with HNP-1, IVIg, or both. Two-tailed paired t-tests were used for comparison of HAdV vs. HAdV + HNP-1 (n = 25, grey). Samples in black (n = 3) were used for analyses between HAdV+ HNP-1 vs. HAdV+ HNP-1 +IVIg.

The receptors for HAdV-C5, -D26, and -B35 partially overlap in human DCs (24): HAdV-C5 predominantly uses DC-SIGN (CD209) (25, 26); HAdV-B35 predominantly uses CD46 (27), and HAdV-D26 uses sialic acid-bearing glycans (28) and CD46 (29). We therefore tested the impact of HNP-1 on capsid uptake using ΔE1 vectors containing reporter genes. Transgene expression was used as a surrogate for endocytosis, cytoplasmic transport, delivery of the capsid to the nuclear pore, and detection of the transgene. We selected doses to have ~30% of DCs expressing the vector-encoded transgene throughout this study. Consistent with earlier reports (25), we found that HNP-1 increased (*p* ≤ 0.02) HAdV-C5 uptake by monocyte-derived DCs (**Figure 1C & D**). Monocyte-derived DCs incubated with HAdV-D26- and -B35-HNP-1 complexes also more readily expressed the transgene (*p* ≤ 0.0001) than each HAdV alone (**Figure 1C & D**). The change in median fluorescence intensity (MFI) and geometric mean of fluorescence intensity (gMFI) corresponded to a 1.6 to 4.1-fold and 1.6 - 3.6-fold increase, respectively, for HAdV-C5, -D26, and -B35. HAdV-D26 showed the lowest change in MFI and gMFI and HAdV-B35 showed the highest (**Figure 1C**). HAdVs preincubated with HNP-1, adding HNP-1 to the cell medium before HAdV, or adding HNP-1 to the medium after HAdV, all increased uptake (**Figure S1B**).

IVIg, which contains a high level of HAdV-C5 neutralizing antibodies, does not prevent HAdV-C5 infection of monocyte-derived DCs but, paradoxically, increases infection (30, 31). To characterize the effects induced by HNP-1, we compared the HAdVs complexed with IVIg or HNP-1. Generally, the impact of HNP-1 was similar to IVIg (**Figure 1D**). We then combined HNP-1 and IVIg with each type to determine if a synergistic effect occurred. However, IVIg did not increase further HNP-1-enhanced transgene expression by HAdV-C5, - D26 or -B35 (**Figure 1D**).

In addition to HNP-1-enhanced uptake by monocyte-derived DCs, we found that HNP-1increased HAdV-C5 and -D26 infection by monocytes (Figure S1C). By contrast, HNP-1 had no additive effect on HAdV-C5- or D26-mediated transgene expression by monocyte-derived Langerhans cells (LCs), whereas it increased HAdV-B35-mediated transgene expression (Figure S1D). Together, these results demonstrated that HNP-1 binds to three HAdV species/types and that its presence tended to increase HAdV infection by primary human APCs.

### HAdV-HNP-1 complexes induce cytokine secretion

Individually, HAdVs and HNPs can promote the activation of DCs by directly engaging PRRs (4, 5, 7). To determine whether there is a synergy between HAdVs and HNP-1, we used a multiplex cytokine array to screen the response by human DCs. As all component of the complexes can individually influences DC maturation, we provide individual baselines to identify the impact of the complex. We used LPS throughout the study as a TLR4 inducer to show the relative range of cytokines and chemokines produced. HAdV-D26 and -B35 induced a greater cytokine response than HAdV-C5 or mock-treated monocyte-derived DCs (**Figure 2A**, 4 columns on the left). We found that HAdV-HNP-1 complexes increased the release primarily in IL-1α and IL-1β (**Figure 2A**, the 2^nd^ group of 4 columns) compared to HNP-1-treated monocyte-derived DCs. When comparing the HAdV vs the HAdV-HNP-1 complexes, the addition of HNP-1 induced an increase in several cytokines (**Figure 2A**, the three pairs of columns on the right). The more marked effect of HNP-1 on HAdV-C5 may be due to the lower effect of HAdV-C5 alone on monocyte-derived DCs (see raw data in **Figure S2A**).

**Figure 2.**
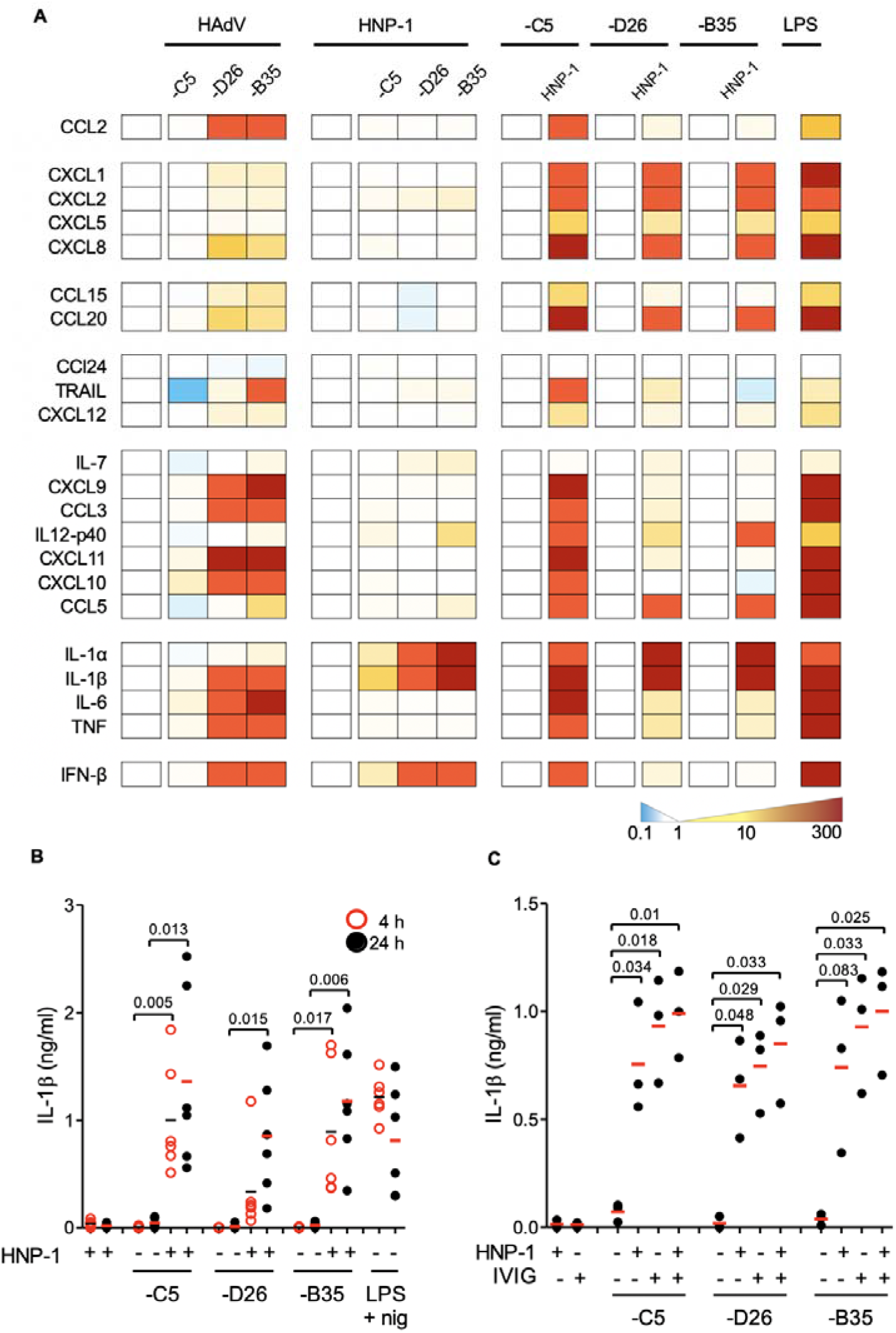
HAdV-HNP-1 complexes induce DCs to release IL-1α and IL-1β. **A)** Monocyte-derived DCs were incubated with HAdV-C5, -D26, or -B35 ± HNP-1, and cytokine secretion in supernatants was assessed by Luminex. To the left of each set of columns is a blanked reference. For the 4 HAdV columns on the left, the reference is mock-treated cells; for the middle HNP-1columns the reference is HNP-1-treated cells; for the “paired” columns on the right (-C5, -D26, and -B35) the reference is HAdV-infected cells compared to HAdV-HNP-1 infected cells. Raw data can be found in (**Figure S2A**). LPS was used as a positive control for TLR4 activation. **B)** Mature IL-1β release induced by HAdV-C5, -D26, and -B35 ± HNP-1 at 4 h (red circles) and 24 h (black dots) post-incubation (n = 5). As above, cells were treated with LPS and nigericin as controls. **C)** DCs were incubated with HAdV-C5, -D26, or -B35 ± HNP-1, IVIg, or HNP-1+IVIg. Mature IL-1β release at 24 h post-incubation (n =3). Statistical analyses were by two-tailed paired t-test unless otherwise noted.

We then quantified the release of mature IL-1β. The HNP-1 + HAdV complexes induced greater IL-1β release than each HAdV alone, which increased from 4 to 24 h post-stimulation (*p* ≤ 0.017 and *p* ≤ 0.013, respectively) (**Figure 2B**). Monocytes, which typically generate low levels of IL-1β, did not release more IL-1β when challenged with HAdV-HNP-1 compared to HNP-1 alone (**Figure S2B**). To characterize the impact of HNP-1 on monocyte-derived DCs, we compared HNP-1 to the effect induced by IVIg-complexed HAdVs. We found that IVIg-complexed HAdV induced modestly more IL-1β release than HNP-1-complexed HAdVs (**Figure 2C**). When IVIg and HNP-1 are combined with the HAdVs, a minor increase was also seen (**Figure 2C**). Using phenotypic (CD86 surface level) and functional (changes in phagocytosis) assays, we found that HNP-1 increased DC maturation when using HAdV-C5 and -D26 (**Figure S2C-D**). Together, these data demonstrated that HNP-1 increased the innate immune response to HAdV-C5 and -D26 and led to functional and phenotypic maturation of monocyte-derived DCs. Notably, these data also mirror the increased efficacy of HAdV infection induced by HNP-1.

### HAdV-HNP-1 complexes induce monocyte-derived DC maturation through TLR4 engagement

As α-defensins are structurally similar to β-defensins, which can activate DCs in a TLR4-dependent manner (32), we explored the possibility that HNP-1-enhanced HAdV transgene expression was due to TLR4 engagement. We used TAK-242, a cell-permeable cyclohexene-carboxylate, to disrupt TLR4 - TIRAP and TRAM interactions, which influence NF-κB activation (33–35). We found that TAK-242 reduced (*p* ≤ 0.019) HNP-1-enhanced uptake of HAdV-C5, -D26 and -B35 (**Figure 3A**). The influence of TAK-242 on IL-1β release (**Figure 3B**) reflected the infection assay: we found a reduction (*p* ≤ 0.045) post-incubation with HAdV-C5 and -B35-HNP-1 challenge, but less prominent for HAdV-D26. Yet, in each case pre-incubation with TAK-242 reduced extracellular IL-1β levels to background levels. Consistent with other reports, TAK-242 reduced (*p* ≤ 0.05) TNF secretion induced by HNP-1-HAdV complexes and LPS (**Figure 3C**). Of note, HNP-1 did not significantly enhance infection of monocytes or Langerhans cells (**Figure S3A & B**), suggesting differences in TLR4 function(s) and/or stoichiometry between these APCs.

**Figure 3.**
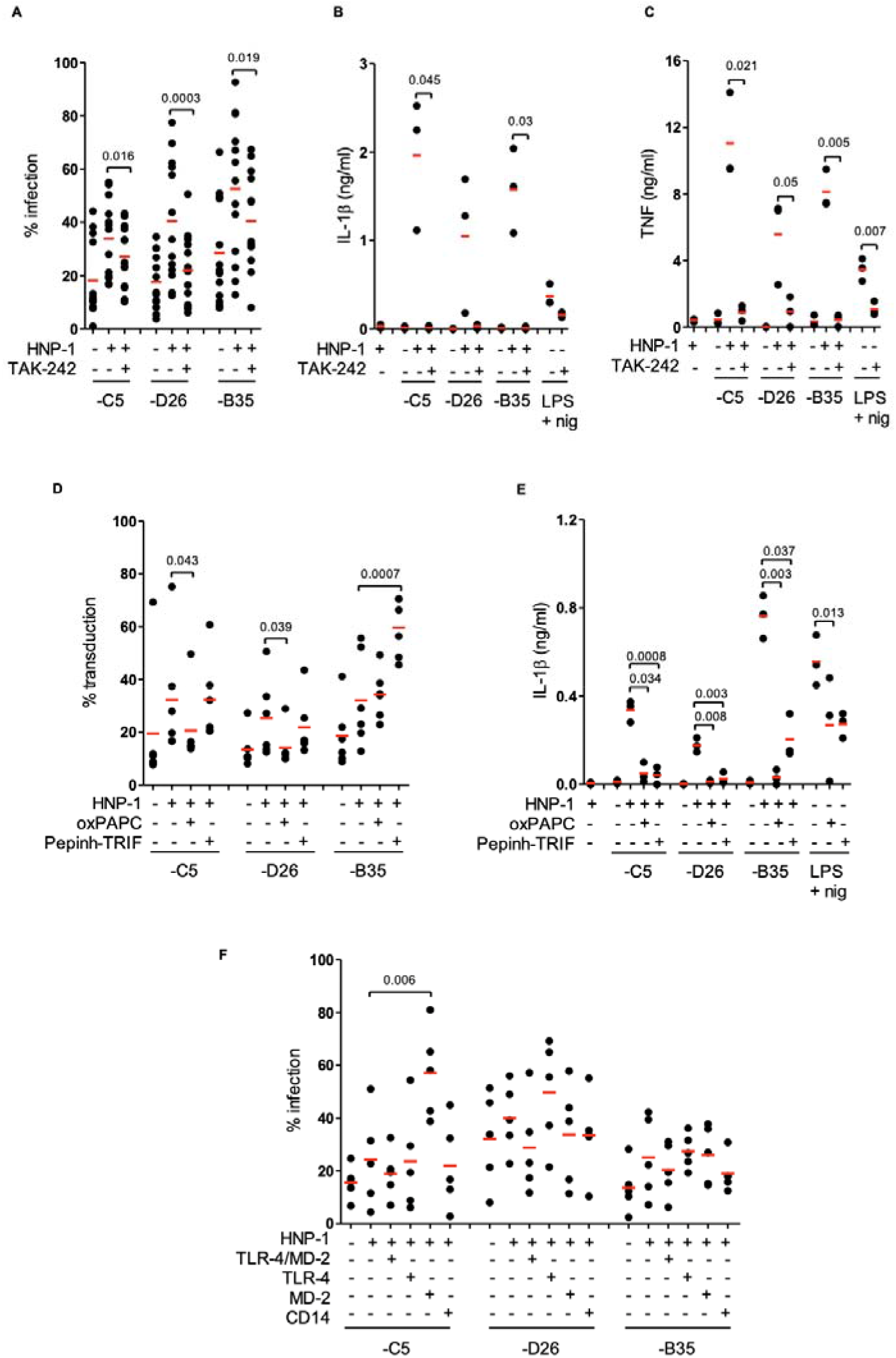
TLR4 engagement and signalling involved in HAdV-HNP-1DC infection. **A)** Monocyte-derived DCs were treated with TAK-242, challenged with HAdV-HNP-1 complexes, and analyzed by flow cytometry (n = 13). **B)** Mature IL-1β release post-TAK-242 treatment (n =3 Student t-test). **C)** TNF levels ± TAK-242 (n = 3). **D)** Percentage of infection following inhibition with oxPAPC or Pepinh-TRIF (n = 6). **E)** IL-1β release following challenge by HAdV-HNP-1 ± oxPAPC or Pepinh-TRIF (n = 3). **F)** HAdV-HNP-1 complexes were challenged with recombinant TLR4, TLR4/MD-2, MD-2 proteins, or with an anti-CD14 antibody. Then, recombinant protein-HAdV ± HNP-1 were added to DCs. Transgene expression was analyzed 24 h post-incubation (n = 5). Statistical analyses were by two-tailed paired t-test unless otherwise noted.

To assess the role of the extracellular TLR4 domain in HNP-1-enhanced infection, we used oxPAPC, which interferes with TLR4 - MD2 interactions. oxPAPC reduced infection by HAdV-C5- or HAdV-D26-HNP-1 complexes (*p*< 0.043), but not HAdV-B35 (Figure 3D). We then used Pepinh-TRIF to inhibit cytoplasmic TLR4 - TRIF interactions and found that Pepinh-TRIF had no effect on HAdV-C5 and -D26, but, surprisingly, increased (*p* < 0.0007) HNP-1-HAdV-B35 infection (Figure 3D). Additionally, oxPAPC and Pepinh-TRIF decreased (*p* < 0.037) IL-1β release induced by HNP-1-HAdVs complexes (Figure 3E). Combined, these data suggested that TLR4 signalling is involved in HNP-1-enhanced infection, and IL-1β and TNF production.

MD-2 can form a complex with TLR4, with the former acting as a co-receptor for exogenous and endogenous ligands (36) or with CD14, for ligand internalization (37). We therefore assayed whether TLR4 engagement is influenced by recombinant TLR4, MD-2, TLR4/MD-2 dimers, or a CD14-blocking antibody. Using this approach, only recombinant MD-2 notably modified HNP-1-enhanced infection by increasing (*p* ≤ 0.006) HAdV-C5 infection. Recombinant TLR4 trended (*p* > 0.05) toward an increase in HAdV-D26 infection (**Figure 3F**). Together, these data are consistent with TLR4 complex engagement and that intracellular TLR4 signalling is involved in the sensing of HNP-1-complexed HAdV-C5, -D26 and -B35.

### HAdV-HNP-1 complexes induce NLRP3 inflammasome formation

Classic and alternative inflammasome activation are associated with IL-1β release. Canonically, there is a 2-step pathway: the initial step induces the transcriptional and translational upregulation of inflammasome components, and the second step is inflammasome formation. To determine whether HAdV-HNP-1engagement of TLR4 induced the transcription of inflammasome components, we used RT-qPCR to examine *NLRP3, CASP1*, and *IL1B* mRNAs. HNP-1-complexed HAdVs induced higher (*p* ≥ 0.012) levels of all three mRNAs compared to HAdVs alone. However, these increases were only higher than HNP-1 alone for ***NLRP3*** for HNP-1-HAdV-B35 (**Figure S4A-C**), suggesting that HNP-1-induced levels were near saturation.

**Figure 4.**
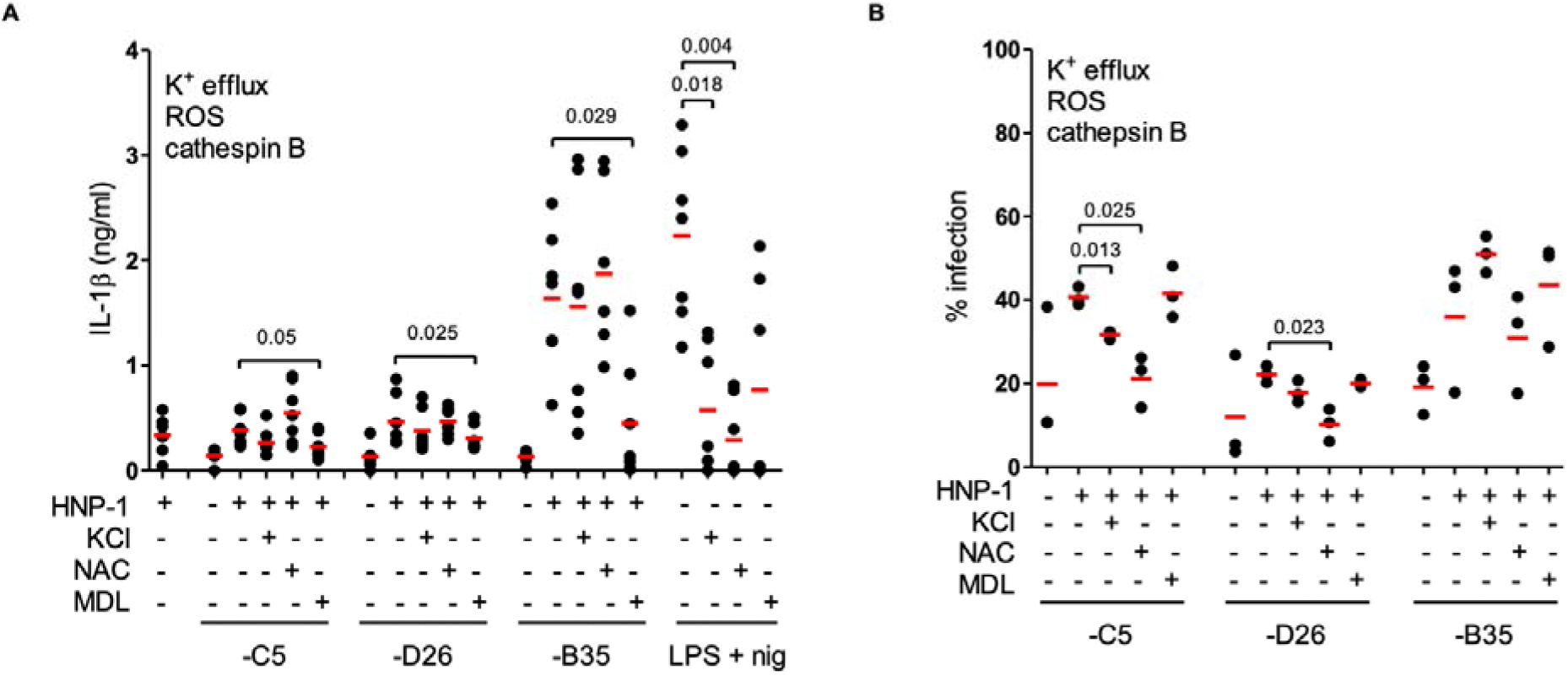
HAdV-HNP-1 complexes induce endosomal rupture. **A)** Monocyte-derived DCs were pre-treated with KCl, NAC, and MDL to inhibit inflammasome formation (n = 6). **B)** The percentage of infected DC following KCl, NAC, and MDL (n = 3) pre-treatment; NAC =N-acetyl-L-cysteine, MDL = MDL 28170. Statistical analyses were by two-tailed paired t-test unless otherwise noted.

To identify the HAdV-HNP-1-associated trigger(s) for inflammasome induction, monocyte-derived DCs w**ere** pre-treated with KCl (to prevent K^+^ efflux), N-acetyl-L-cysteine (NAC) (a ROS scavenger), and MDL (to inhibit endosomal proteases-mediated activation of the NLRP3 inflammasome after endosome lysis) (12). LPS in the presence or absence of the NLRP3 inducer nigericin was used as a control. We found that only the addition of extracellular MDL significantly (*p* < 0.05) decreased the release of IL-1β for HAdV-C5, -D26, and -B35 (**Figure 4A**). Adding K^+^ or NAC also modified HNP-1-enhanced infection by HAdV-C5 and -D26 (**Figure 4B**). Of note, following receptor-mediated endocytosis, HAdVs escape from endosomal compartments via the endosomolytic activity of protein VI (38). Therefore, these data are consistent with the idea that endosomal rupture caused by HNP-1-enhanced uptake and cathepsin B release into the cytosol, which in turn induces inflammasome induction and IL-1β release.

We then used small molecule inhibitors to dissect the crosstalk between TLR4-induced signalling and NLRP3 induction in the presence of HNP-1-HAdV complexes. DCs pre-incubated with R406 (blocking MyD88-Syk interactions and interrupting the signalling between TLR4 engagement and NLRP3 inflammasome regulation), or Bay 11-7082 (inhibitor of NLRP3 (39)), IL-1β release was reduced (*p* ≤ 0.038) in response to HNP-1 complexes in all cases barring R406 along with HNP-1 and HAdV-D26 (**Figure 5A**). Bay11-7082 and R406 did not significant impact on HNP-1-enhanced infection (except for HAdV-B35) (**Figure 5B**). MCC950, an NLRP3 inhibitor, also inhibited IL-1β release **(Figure 5C)**.

**Figure 5.**
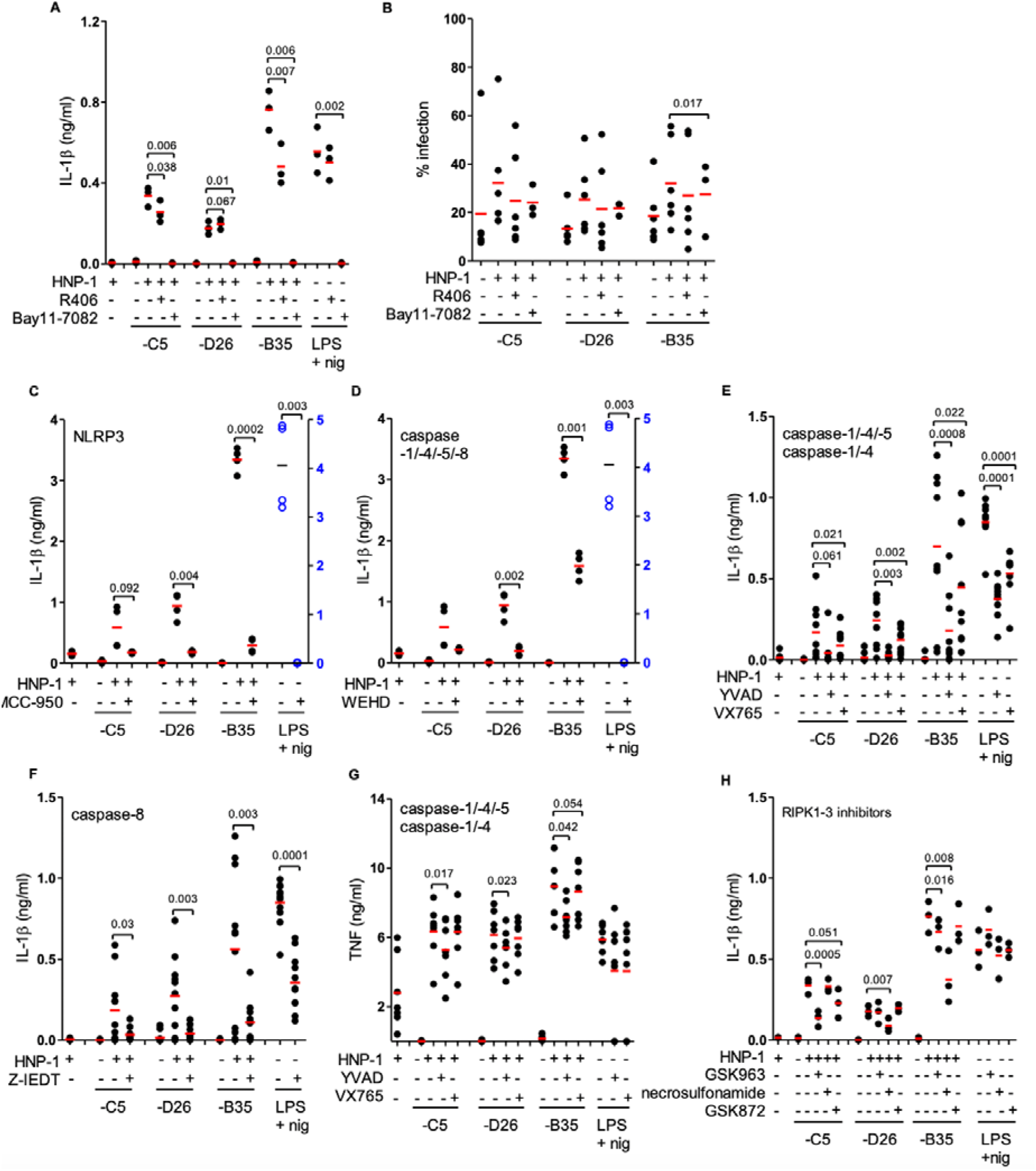
NLRP3 is linked to IL-1β release. **A)** Monocyte-derived DCs were treated with R406 or Bay11-7082 prior to challenge with HAdVs ± HNP-1. LPS/nigericin was used as a control. **B)** DCs were treated with R406 or Bay11-7082 before challenge with HAdVs ± HNP-1. The percentage of DCs expressing the reporter gene. Impact of **C)** MCC-950; **D)** WEHD; **E)** YVAD and VX765; **F)** Z-IEDT on IL-1β release; **G)** Impact of YVAD and VX765 on TNF release. **H)** IL-1β release in response to HAdV-HNP-1 complexes in DCs pre-treated with RIPK1-RIPK3 pathway inhibitors GSK963, necrosulfonamide, or GSK872. (n ≥ 3). Statistical analyses were by two-tailed paired t-test unless otherwise noted.

We then inhibited selected caspases using VX765 (caspase-1 and −4), Z-IETD (capsase-8), WEHD (caspase-1, −4, −5 and −8), and Z-YVAD-FMK (caspase-1, −4 and −5). We found that all the inhibitors reduced IL-1β release to some extent (**Figure 5D-G**). Importantly, the caspase inhibitors had no consistent effect on HNP-1-enhanced infection (**Figure S5A**). In the alternative inflammasome pathway, activation of the NLRP3 inflammasome depends on signalling through TLR4 and the kinase RIPK1, whereas the involvement of kinases RIPK3 and MLKL downstream of RIPK1 is not required (14). Consequently, we used GSK963 (RIPK1 inhibitor), GSK872 (RIPK3 inhibitor), and necrosulfonamide (MLKL inhibitor) to investigate whether the alternative inflammasome pathway was involved. Again, the inhibition of the three kinases had no significant effect on infection (**Figure S5B**). By contrast, GSK963 and necrosulfonamide reduced (*p* ≤ 0.016 and *p* ≤ 0.007, respectively) IL-1β release following HNP-1-enhanced infection of HAdV-C5 and -B35 and by necrosulfonamide during HNP-1-enhanced infection of HAdV-D26 and -B35 (*p* ≤ 0.007) (**Figure 5H**, and **S5C** for analyses broken down by HAdV type).

We concluded that HAdV-HNP-1 complexes induced NLRP3 inflammasome formation and IL-1β release via the activation of the TLR4 pathway. Of note though, the pathway does not fully recapitulate all steps of the alternative inflammasome pathway. Furthermore, the cellular factors involved in HNP-1-enhanced HAdV infection are upstream of NLRP3, caspases and RIPK1, RIPK3 and MLKL kinases.

### IL-1β release and membrane integrity

These data incited us to investigate the activation of the alternative inflammasome pathway (14). To determine whether HAdV-HNP-1 complexes induced the release cytoplasmic proteins, we quantified extracellular activity of L-lactate dehydrogenase (LDH) at 4 h post-incubation. We found no notable (*p* > 0.05) increase in any condition (**Figure 6A**). To determine whether HAdV-HNP-1 complexes impacted membrane integrity, we added 7-AAD and quantified its uptake by DCs. We found that the percentage of 7-AAD^+^ cells induced by HAdV-HNP-1 complexes was greater (*p* ≤ 0.044) than HNP-1- or HAdV-challenged cells (**Figure 6B**). Additionally, when HNP-1 was added to HAdV-IVIg complexes, a modest increase in the percentage of 7-AAD^+^ cells was detected (**Figure 6C**). Therefore, within the time frame of our assays, holes large enough to allow small molecule entry were created, but LDH was not released into the medium, suggesting that pyroptosis was aborted.

**FIGURE 6.**
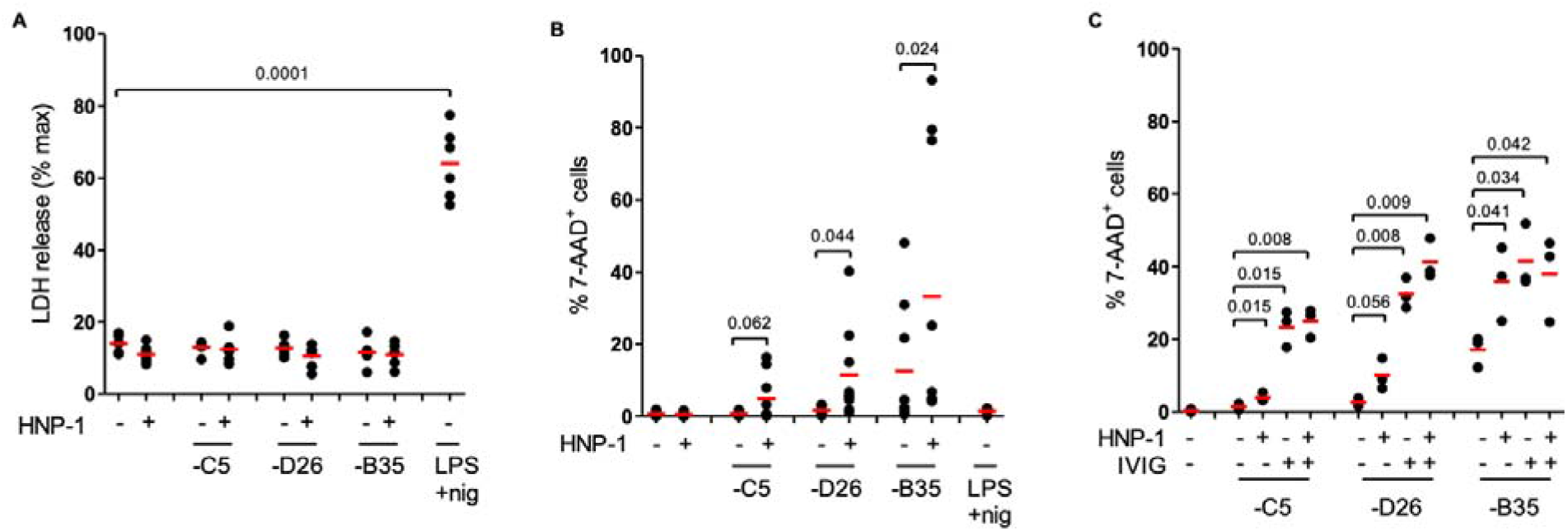
IL-1β release without loss of membrane integrity. **A)** Monocyte-derived DCs were incubated with HAdV-C5, -D26, or -B35 ± HNP-1, or LPS/nigericin. Extracellular LDH activity was quantified at 4 h post-incubation (n = 6). **B)** 7-AAD uptake was analyzed 24 h post-incubation (n = 9). **C)** DCs were incubated with HAdV-C5, -D26, or -B35 ± HNP-1, IVIg or HNP-1 + IVIg. 7-AAD uptake was quantified 24 h post-incubation (n = 3). Statistical analyses were by two-tailed paired t-test unless otherwise noted.

## Discussion

Here, we demonstrate how an α-defensin promotes innate immune sensing of HAdVs and maturation in human DCs. We used biochemical, pharmaceutical, cellular, and molecular approaches to address HAdV-HNP-1 inflammasome formation/activation in primary human APCs. Our data demonstrate that HNP-1 binds to the HAdV capsids, enhances the infection of human phagocytes, and engages TLR4 to induce pro-inflammatory cytokines via the NLRP3 inflammasome.

Based on their efficacy as vaccines and differences in receptor use, it was not surprising to observe similarities and notable differences between the HAdVs assayed in this study. Notably, HAdV-B35 also had the strongest inflammatory response even though it was used at the lowest dose (1,000 pp/cell), which may be attributed to the engagement of CD46 and the TLR4-NLRP3 axis described here. It is worth noting that CD46 engagement can induce the formation of the NLRP3 inflammasome in human CD4^+^ cells (40). We speculate that the individual attachment receptors for HAdV-C5 (CAR and DC-SIGN), HAdV-D26 and HAdV-B35 (CD46) and their stoichiometry are involved in the differences reported here. Notably, CAR is not expressed on most human phagocytes, and crosstalk between TLR4 signalling with DC-SIGN or CD46, respectively, to induce pro-inflammatory responses have been described and may account for the differences seen in our study (41–43). Of note, we did not directly address the differences in viral DNA entering the cells and acting as a PAMP for TLR9 engagement (30).

Smith and Nemerow showed that α- and β-defensins inhibit HAdV infection of epithelial cells (8). HNP-1, similar to lactoferrin, enhances infection of HAdV by human monocytes, DCs, and to some extent LCs. This may indicate that defensins redirect HAdVs from infection of epithelial cells to innate immune cells to initiate an antiviral response. Because HNP-1 can stimulate monocyte, DC, and T-cell recruitment (32, 44–46) this synergistic effect likely plays a significant role in the downstream adaptive response. By exploring further these early events, we address the mechanisms by which an α-defensin modifies innate and adaptive immune responses to HAdVs.

Like lactoferrin, HNP-1-enhanced HAdV infection induced a pathway similar to the alternative NLRP3 inflammasome pathway (11), which is characterized by the activation of TLR4-TRIF-RIPK1-FADD-CASP8 cascade and the absence of transcriptional priming step, K^+^ efflux, and pyroptosome formation (14). While the key features are well aligned between the alternative pathway and our data, it is notable that the inhibition of RIPK1 only partially abrogated IL-1β secretion induced by HNP-1, and that MLKL kinase may be involved in the response to HAdV-D26 and -B35. These differences may be due to the cell types used (i.e. DC versus monocytes) (14), which express different levels of CD14 that affect TLR4 stoichiometry and signalling. More importantly, LPS and HAdVs substantially differ in size, molecular composition, and functional factors that affect intracellular processing and cytosolic PAMP detection. TLR4-mediated endocytosis of LPS involves sequential interaction of CD14 and MD2 with LPS and depends on the homodimerization of TLR4. The shape and size (~950 Å icosahedron) of the HAdV-HNP-1 complex allow HNP-1 to bind the capsid and induces TLR4 dimerization or even multiple dimers directly. Quintessential TLR4 agonists do not have complex intracellular processing. However, HAdVs traffic through endosomal compartments until the pH drops sufficiently to allow capsid escape into the cytosol. When a HAdV capsid is covered with neutralizing antibodies, lysosomal degradation of the capsids is more efficient, the viral genome is partially exposed, and the DNA is then detected via TLR9 (30, 47). Crosstalk between TLRs and synergistic effects on IL-1β secretion has been previously reported for human DC (48) *in vitro* and in mice (49, 50). Furthermore, hierarchical dependency of cell surface receptors (TLR4) and intracellular PRRs in phagocytes can affect the intracellular processing of the “ligand” (51). Notably, also TLR4 engagement channels its ligands, in a TRIF-dependent manner, into specialized intracellular compartments (52) and intracellular trafficking of a PAMP through endosomal and lysosomal compartments and the cytosol can lead to differential signalling and the production of type I IFN, TNF and IL-1β (53, 54). In addition, HNP-1 increases the sensitivity of human plasmacytoid DCs against TLR9 ligands and thus, promotes the induction of NF-κB and IRF1 response genes (55, 56). The range of our assays indicates that TLR4 signalling though MyD88 is required for HNP-1 enhanced HAdV infection while signalling through MyD88 and TRIF is required for the induction of the NLRP3 inflammasome. We did not investigate how HAdV processing occurs downstream of TLR4-mediated uptake, however, it is possible that the HNP-1-dependent increase of transgene expression coincides with increased amounts of intracellular viral DNA and disrupted intracellular vesicles. It is possible that the higher levels of viral DNA stimulate TLR9 or other PRRs and account for the differences seen between the alternative inflammasome pathway and our data. Of note, our data also indicate that cathepsin B release from endosomal compartments concomitant with HAdV endosomal escape induces NLRP3 induction and IL-1β secretion, which may further add to the differences seen by us and Gaidt *et al*. (14).

Here, HAdV-HNP-1 induced signals are received immediately prior to NLRP3 engagement. We hypothesize that the coordination of priming and de-ubiquitination of NLRP3 (57), as well as TNF survival signals (58), favour an adaptive immune response. Inflammasome activation is crucial for inducing cellular and humoral immune responses. Recently, moderate NLRP3 activation in human conventional type 2 DCs induced a hyperactivated phenotype characterized by secretion of IL-1β and IL-12 family cytokines in the absence of pyroptosis and followed by strong induction of Th1 and Th17 responses (59). Of note, HNP-1 also enhanced HAdV-induced IL12-p40 secretion in our assays (**Figure. 2A**). We speculate that the involvement of the HAdV-HNP-1-TLR4-NLRP3 axis may help drive the T-cell responses towards a Th1 phenotype. Recent clinical trials indicate that Ad26.COV2.S, a HAdV-D26-based vaccine, induces potent Th1 responses against the SARS-CoV-2 spike (60, 61). It is conceivable that HNP-1 acts as a natural adjuvant and increases the innate response and ultimately the breadth of the adaptive response to intracellular pathogens. In addition, controlling this inflammatory response may be due to the production of IL-1α, which promotes cell survival (62).

In conclusion, we examined the multi-layered interface of human innate and adaptive responses to HAdV pathogenesis and HAdV-based vaccines.

## Materials and Methods

### Cells and culture conditions

Blood samples were purchased from the regional blood bank (EFS, Montpellier, France). An internal review board approved their use. MoDCs were generated from freshly isolated or frozen CD14^+^ monocytes (30). DC stimulation was performed 6 d post-isolation. Monocyte-derived LC were generated using 200 ng/ml GM-CSF and 10 ng/ml TGF-β. 911 and 293 E4-pIX cells were grown in Dulbecco’s modified Eagle medium (DMEM) and minimum essential medium (MEMα) with Earle’s salts, L-glutamine supplemented with 10% foetal bovine serum (FBS).

### Replication-defective vectors

HAdV-C5 contained a GFP expression cassette (63). HAdV-D26 contained a GFP-luciferase fusion expression cassette (17). HAdV-B35 contained a YFP expression cassette (8). Vectors were propagated in 911 or 293 E4-pIX cells and purified by density gradients (63). For infection assays we used HAdV-C5 (5,000 physical particles (pp)/cell), -D26 (20,000 pp/cell) or -B35 (1,000 pp/cell) (red). The samples were collected 24 h later, prepared for flow cytometry and 25,000 events were acquired/sample.

### DC stimulation with HAdV-HNP-1 complexes

DCs (4 x 10^5^ in 400 μl of complete medium) were incubated with HAdV-C5, -D26 or -B35 (0.1 to 2 x 10^4^ physical particles (pp)/cell). HAdV-HNP-1 complexes were generated by incubating the HAdVs with 1.40 μg HNP-1 (Sigma-Aldrich) (64, 65) for 30 min at room temperature. When noted, cells were complexed with IVIg (human IgG pooled from between 5,000 and 50,000 donors/batch) (Baxter SAS). Cells were incubated with HAdV-HNP-1 for 4 h, then washed and incubated for 24 h. LPS (Sigma-Aldrich) and NLRP3 inflammasome inducer nigericin (InvivoGen) were used at 100 ng/ml and 10 μM, respectively. Inhibitor concentrations, TAK-242 (Merck Millipore) at 1 μg/ml, oxPAPC (InvivoGen) at 30 μg/ml, TRIF inhibitory peptide (InvivoGen) at 25 μM, Syk inhibitor R406 (InvivoGen) at 5 μM, KCl (Sigma-Aldrich) at 40 mM, N-acetyl-L-cysteine (Sigma-Aldrich) at 2 mM, MDL 28170 (Tocris Bioscience) at 0.1 mM, MCC-950/CP-456773 (Sigma-Aldrich) at 10 μM, Bay11-7082 (Sigma-Aldrich) at 10 μM, WEHD (Santa Cruz) and YVAD (InvivoGen) at 20 μM, VX765 (InvivoGen) at 10 μM, caspase-8 inhibitor Z-IEDT at 20 μM, GSK963 (Sigma-Aldrich) at 3 μM, GSK872 (Merck Millipore) at 3 μM and necrosulfonamide (R&D systems) at 1 μM. TLR4/MD-2, TLR4 (R&D Systems), MD-2 (PeproTech) recombinant protein and anti-CD14 antibody (Beckman) were used at 20 μg/ml. Inhibitors were added on cells and recombinant proteins or antibody were added on HAdV-HNP-1 complex 1 h before stimulation.

### Surface plasmon resonance analyses

SPR analyses were performed on a BIAcore 3000 apparatus in HBS-EP buffer (10 mM HEPES, 150 mM NaCl, 3 mM EDTA, and 0.005% (v/v) polysorbate 20, pH 7.4). HAdV-C5, -D26 and -B35 diluted in acetate buffer at pH 4 were immobilized on three flow cells of a CM5 sensor chip by amine coupling. Immobilization levels were between 3,500 and 4,000 RU. A Blank flow cell was used as a control. HNP-1 was injected at 100 nM on the 4 flow cells simultaneously. T determine the KD, 6.25 - 200 nM of HNP-1 was injected at 30 μl/min during 180s association and 600s dissociation with running buffer. Regeneration was performed with Gly-HCl pH 1.7 pulses. The kinetic constants were assessed from the sensorgrams after double-blank subtraction with BIAevaluation software 3.2 (GE Healthcare) (GE Healthcare) using a 2-state fitting model. All experiments were repeated at least twice for each vector on a freshly coated flow cells.

### Flow cytometry

GFP or YFP expression from the HAdV-C5, -B35, -D26 vectors was assayed by flow cytometry. CD86 surface expression level was assessed with an anti-CD86 antibody (clone 2331, APC, BD Biosciences). For dextran uptake assays, we used 1 mg/ml for 30 min at 37°C, or 4°C for negative control) (dextran TRITC, Sigma). Cell membrane integrity was assessed by collecting cells by centrifugation 800 x g, the cell pellets were re-suspended in PBS, 2% FBS, 1 mM EDTA, 7-aminoactinomycin D (7-AAD) (Becton-Dickinson Pharmigen) and analyzed on a FACS Canto II (Becton-Dickinson Pharmigen) or NovoCyte (ACEA Biosciences) flow cytometer.

### Cytokine secretion & LDH release

Supernatants were collected 4 or 24 h post-challenge and the levels of TNF and mature IL-1β were quantified by ELISA using OptEIA human TNF ELISA Set (BD Biosciences) and human IL-1β/IL-1F2 DuoSet ELISA (R&D systems). Twenty-two cytokines were detected using Bio-plex human chemokine, cytokine kit (Bio-Rad). LDH release was quantified using an LDH Cytotoxicity Assay Kit (Thermo scientific) and as previously described (11).

### Quantification of *NLRP3, CASP1* and *IL1B* mRNAs

mRNAs levels were analyzed using qRT-PCR as previously described (11).

## Acknowledgments

We thank Eric Weaver and Andre Lieber for HAdV-D26 and -B35 vectors. We acknowledge the MRI (ANR-10-INBS-04). This work benefited from support from Université de Montpellier (EJK), Ph.D. fellowships from the French Minister of Education (OP) and the Vietnamese Minister of Education (TTPT), TransVacII, and EpiGenMed, an “Investissements d’avenir” program (EJK). EJK is an Inserm Fellow.

The funders played no role in study design, data collection and analysis, decision to publish, or preparation of the manuscript.

## Author contributions

Study design & conception: KE, EJK

Project direction: EJK

Performed experiments; CC, KE, HT, TTPT, OP, CH

Analysed data: all authors

Wrote the manuscript: KE, CC & EJK

Secured funding: EJK

The authors declare that there are no competing interests.

## Data Availability

All data generated or analyzed during this study are included in this published article (and its supplementary information files).

## Supplementary data

**Figure S1).**
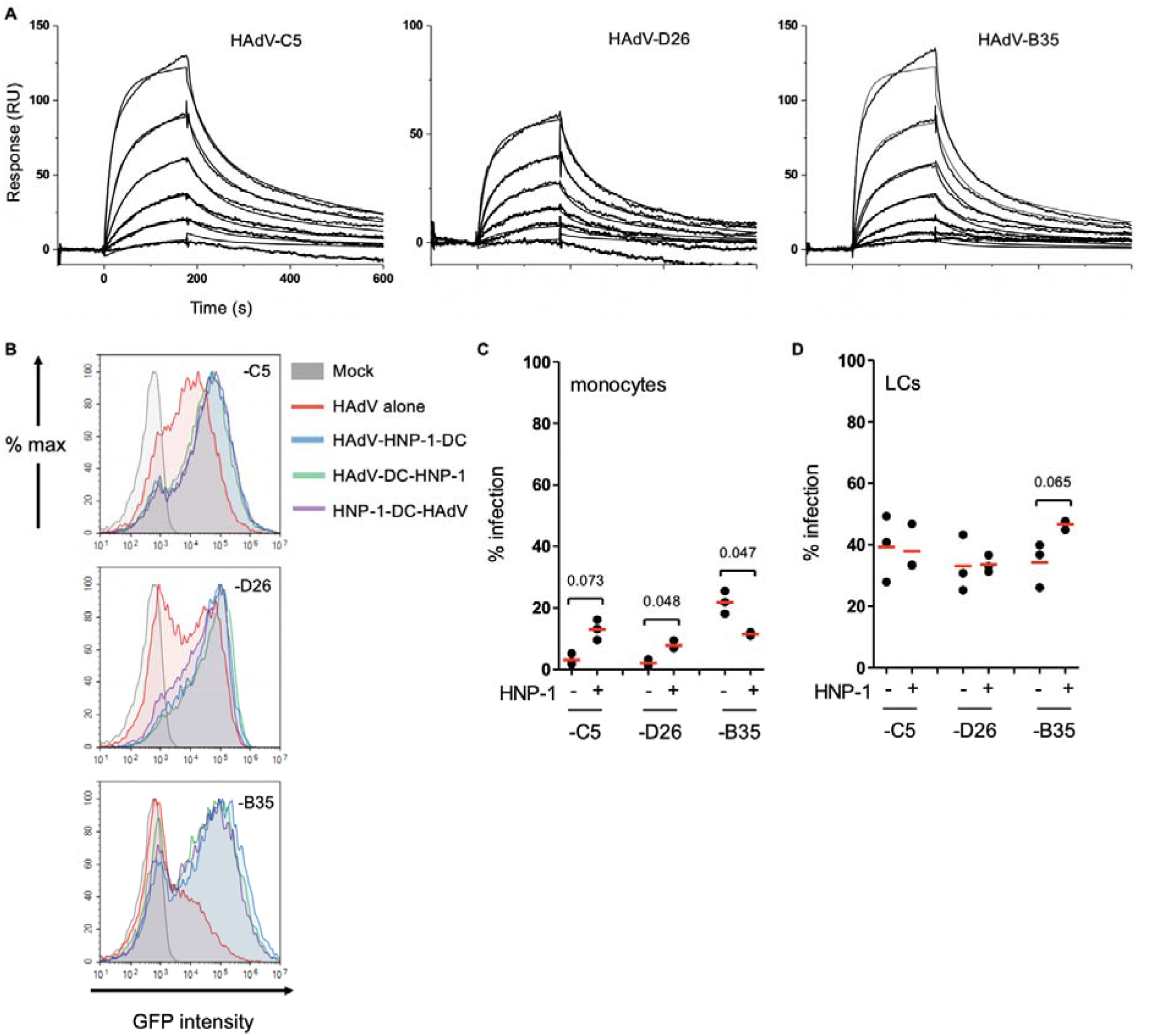
Analyses of HNP-1 binding to HAdV-C5, -D26 and -B35 by SPR. **A)** HAdV-C5, -D26, and -B35 were covalently coupled to a CM5 sensor chip and escalating doses of HNP-1 (6.25-200 nM) for KD determination. Depicted are overlaid sensorgrams (RU = resonance units). **B)** Representative flow cytometry profiles of cells incubated with HAdVs ± HNP-1. DCs were mock-treated (grey), incubated with HAdV - C5, -D26 and -B35 alone (red), with HNP-1 complexed with HAdV (blue), with HAdV for 30 min and then HNP-1 (green) or with HNP-1 for 30 min and then HAdV (purple). Fluorescence was analyzed 24 h post-incubation. **C)** monocytes and **D)** LCs were incubated with HAdVs ± HNP-1 and GFP/YFP expression was analyzed 24 h post-incubation (n = 3, statistical analyses by two-tailed paired t-test).

**Figure S2).**
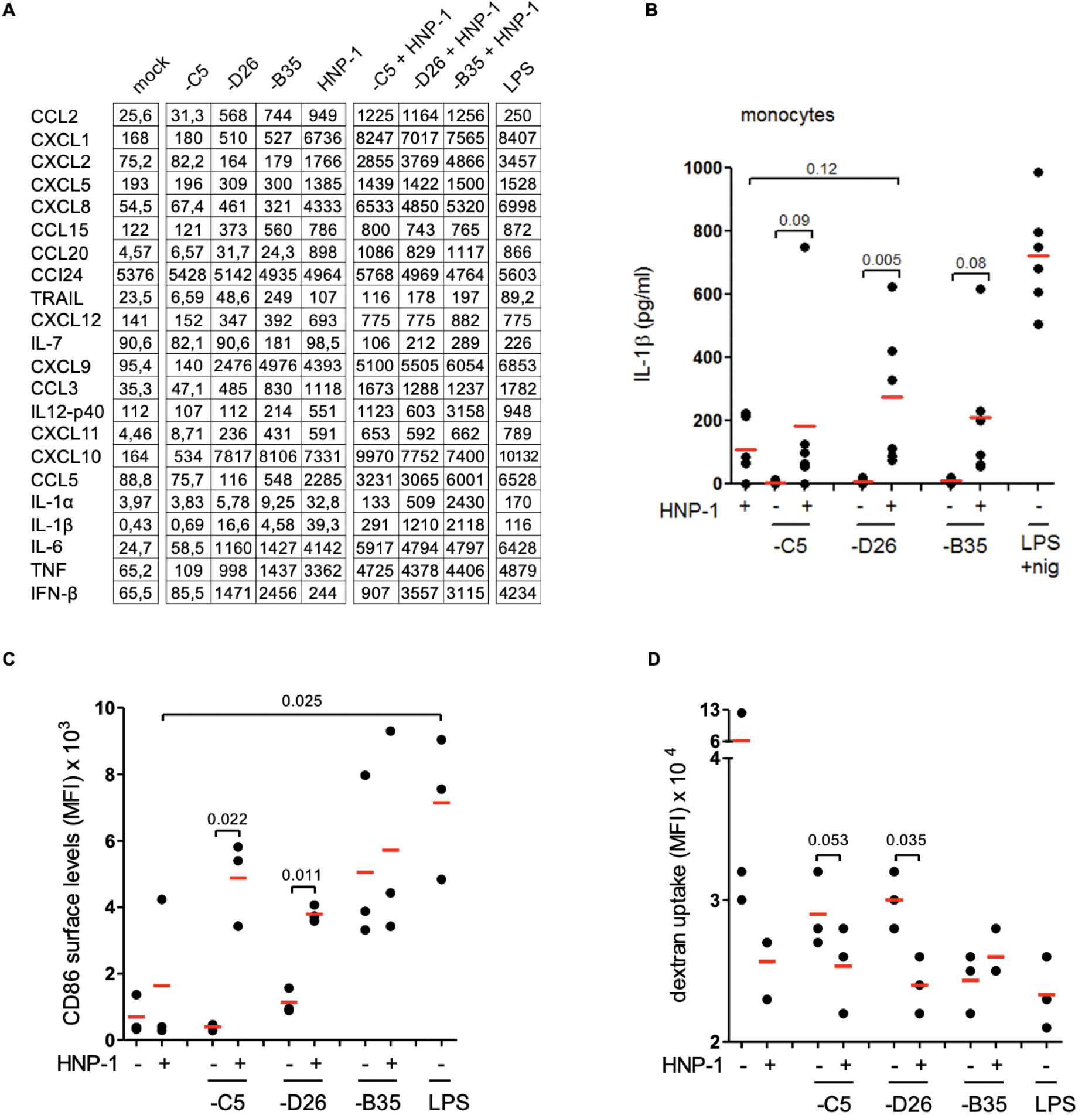
HAdV-HNP-1 induced cytokine secretion and DC maturation. **A)** Raw data of DCs ± HAdVs ± HNP-1, plus controls. Data indicated total protein level (ng). **B)** Freshly isolate human CD14^+^ monocytes were incubated with HAdV-C5-, -D26, or -B35 ± HNP-1. The supernatants (24 h post-incubation) were used to quantify IL-1β release (n=6, statistical analyses by Student’s t-test). There was no significant difference between HNP-1 vs. HNP-1+HAdVs. **C)** DCs were incubated with HAdV-HNP-1 complexes for 24 h. Phenotypic maturation was assessed by CD86 surface levels expression and functionally. **D)** DCs were incubated with HAdV-HNP-1 complexes and phagocytosis was quantified by fluorescent dextran uptake. Cells were incubated at 4°C or 37°C for 30 min with 1 mg/ml TRITC-labelled dextran, washed with PBS, fixed with 4% PFA and analyzed by flow cytometry (lower fluorescence = lower phagocytosis = greater maturation) (n =3, statistical analyses by two-tailed paired t-test).

**Figure S3:**
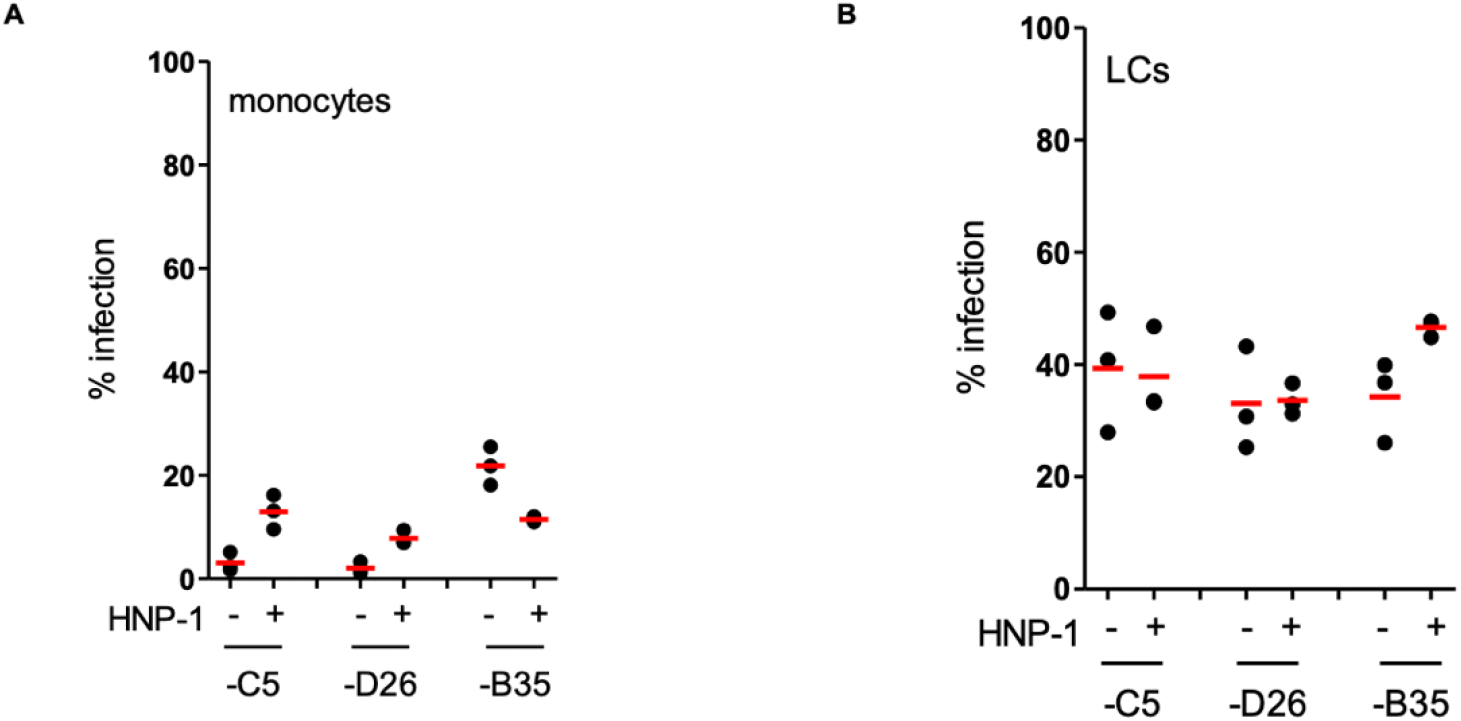
HNP-1-mediated HAdV infection of monocytes and monocyte-derived LCs. **A)** HNP-1-mediated HAdV infection of monocytes was analyzed 24 h post-incubation by flow cytometry (n = 3, statistical analyses by two-tailed paired t-test). **A-B)** HNP-1-mediated HAdV infection of monocyte-derived LCs was analyzed 24 h post-incubation by flow cytometry (n = 3, statistical analyses by two-tailed paired t-test).

**Figure S4:**
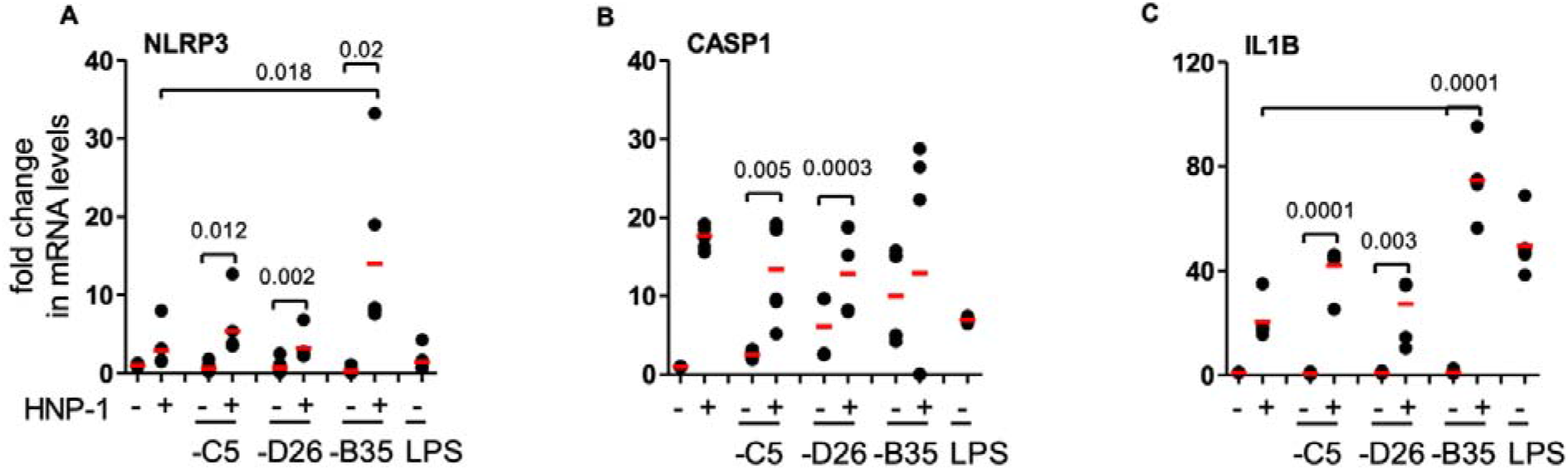
HNP-1 induce transcription of NLRP3 inflammasome. **A-C)** mRNA levels of inflammasome components (encoded by *NLRP3, CASP1*, and *IL1B*) were analyzed using qRT-PCR 4 h post-incubation. Total RNA was isolated from DCs and cDNA samples were analyzed in triplicate. *GAPDH* mRNA was used as an internal standard.

**Figure S5).**
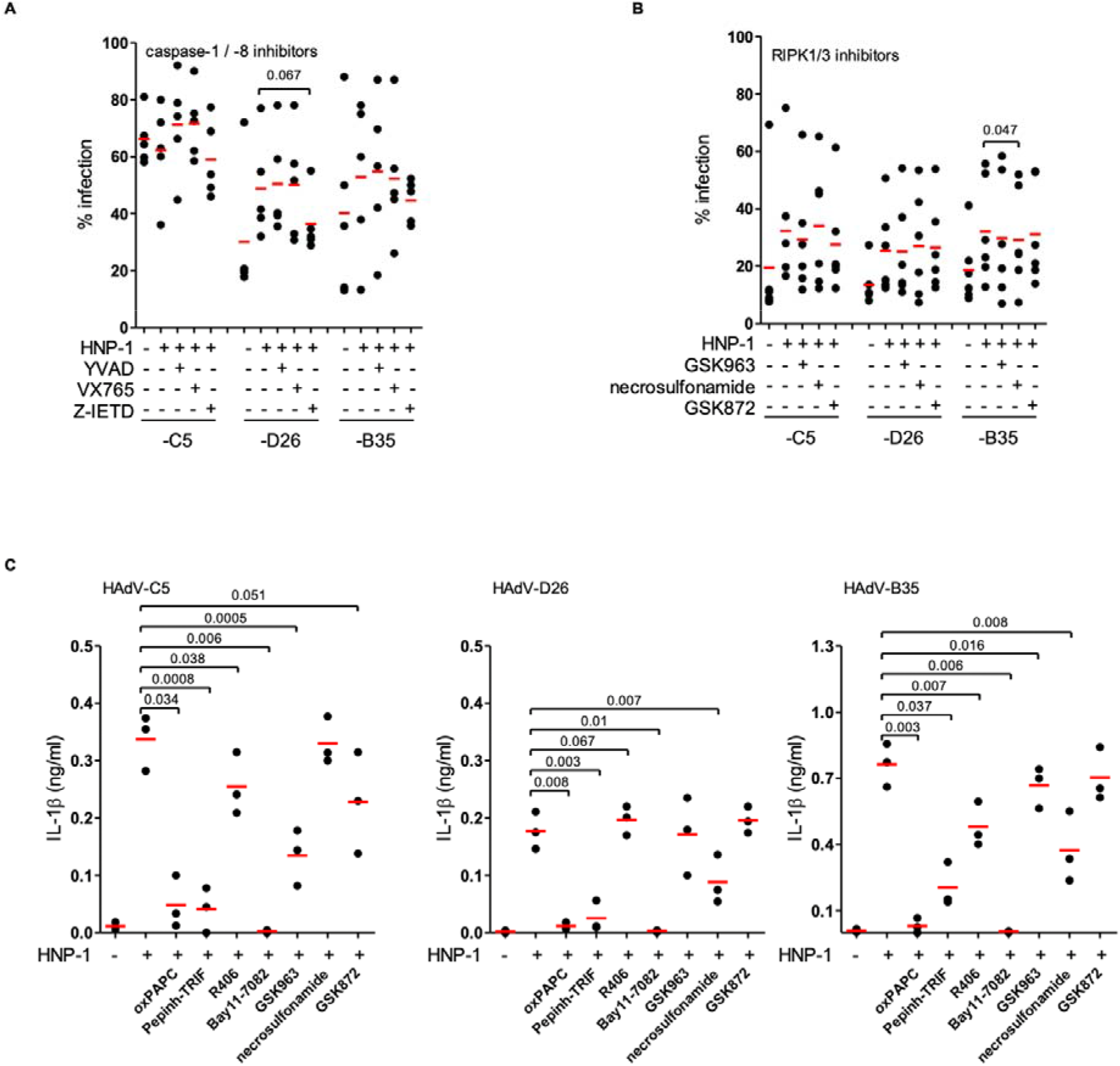
Impact of TLR4, RIPK1, RIPK3 inhibition on HNP-1-HAdV complex induced IL-1β release broken down by HAdV type. **A)** Impact of YVAD, VX765 and Z-IETD (caspase-1, −4, −5, and 8 inhibitors) on infection. **B)** Impact of GSK963, necrosulfonamide, and GSK872 (RIPK1/3 inhibitors) on infection. **C)** Combined analyses broken down by HAdV species/type.

